# Inhibition of DNAJ-HSP70 interaction improves strength in muscular dystrophy

**DOI:** 10.1101/2020.01.03.893149

**Authors:** Rocio Bengoechea, Andrew Findlay, Ankan Bhadra, Hao Shao, Kevin Stein, Sara Pittman, Jil Daw, Jason E. Gestwicki, Heather True, Conrad C. Weihl

## Abstract

Dominant mutations in the HSP70 co-chaperone DNAJB6 cause a late onset muscle disease termed limb girdle muscular dystrophy type 1D (LGMD1D), which is characterized by protein aggregation and vacuolar myopathology. Disease mutations reside within the G/F domain of DNAJB6, but the molecular mechanisms underlying dysfunction are not well understood. Using yeast, cell culture, and mouse models of LGMD1D, we find that the toxicity associated with disease-associated DNAJB6 requires its interaction with HSP70, and that abrogating this interaction genetically or with small molecules is protective. In skeletal muscle, DNAJB6 localizes to the Z-disc with HSP70. Whereas HSP70 normally diffuses rapidly between the Z-disc and sarcoplasm, the rate of HSP70’s diffusion in LGMD1D mouse muscle is diminished likely because it has an unusual affinity for the Z-disc and mutant DNAJB6. Treating LGMD1D mice with a small molecule inhibitor of the DNAJ-HSP70 complex re-mobilizes HSP70, improves strength and corrects myopathology. These data support a model in which LGMD1D mutations in DNAJB6 are a gain-of-function disease that is, counter-intuitively, mediated via HSP70 binding. Thus, therapeutic approaches targeting HSP70:DNAJB6 may be effective in treating this inherited muscular dystrophy.

## Introduction

Limb-girdle muscular dystrophies (LGMD) are a family of hereditary muscle diseases that are inherited in an autosomal recessive or dominant manner [1]. Most recessively inherited LGMDs are postulated to be caused by a loss-of-function mechanism, because the protein is absent in patient muscle tissue and many mutations lead to premature stops or frameshifts. However, the mechanisms of dominantly inherited LGMDs are less clear: they may be due to haploinsufficiency, a gain of new toxic function or a dominant interaction leading to protein dysfunction.

One common pathologic feature seen in some dominantly inherited myopathies, including LGMD1D, myofibrillar myopathy and inclusion body myopathy, are protein aggregates and myofiber vacuolation [2]. The inclusions in LGMD1D muscle are heterogeneous and contain myofibrillar proteins, such as desmin and myotilin [3, 4]. Additionally, they contain proteins that aggregate in neurodegenerative disorders, such as TDP-43 and SQSTM1 [3, 4]. Together, these features suggest that a subset of inherited myopathies might involve an impairment in protein quality control. Indeed, one family of muscle diseases with myofibrillar disorganization (*e.g*. myofibrillar myopathies) are caused by missense mutations in Z-disc associated structural proteins, such as desmin, myotilin or filamin-C that lead to their aggregation [2]. Likewise, myofibrillar myopathies are also associated with mutations in molecular chaperones, such as BAG3, HSPB5 and DNAJB6 [2, 5, 6]. These “chaperonopathies” are autosomal dominantly inherited and lead to the accumulation and aggregation of Z-disc proteins, such as desmin and filamin-C. Moreover, several chaperones are known to be necessary for sarcomere development and maintenance [7]. Our aim was to better understand the mechanisms associated with dominantly inherited LGMD1D caused by mutations in DNAJB6 in order to gain insight into the relationship between protein quality control and myopathy.

DNAJB6 is a ubiquitously expressed HSP40 co-chaperone that facilitates HSP70 functionality via its J domain [8]. DNAJB6, like other DNAJB family members, has a canonical J domain, a C-terminal dimerization motif and a glycine- and phenylalanine-rich region (termed a G/F domain) [8]. Interestingly, all reported LGMD1D mutations reside within the G/F domain [3, 4, 9-11]. DNAJB6 has two isoforms, DNAJB6a and DNAJB6b, which differ in the length of their C-terminal region and in their cellular localization. DNAJB6a is longer and localizes to the nucleus; whereas DNAJB6b is shorter and cytoplasmic [12]. Although both isoforms share the G/F domain containing LGMD1D disease mutations, it is thought that DNAJB6b is the pathogenic isoform [3, 12]. Patients with LGMD1D have insidious onset of weakness in the second to sixth decade of life that affects their ability to ambulate [3, 4, 9, 10, 13]. The muscle tissue of LGMD1D patients is characteristically myopathic, with myofibrillar protein aggregates and rimmed vacuoles [3, 4]. Myoblasts lacking DNAJB6 accumulate the Z-disc associated intermediate filament desmin, suggesting that these phenotypes might involve poor quality control of Z-disc associated proteins [14]. Consistent with this idea, the skeletal muscle from transgenic mice expressing the F93L mutation in DNAJB6b recapitulated features of LGMD1D with muscle weakness, atrophy, myofibrillar disorganization and insoluble desmin aggregates [12].

Yeast has been a powerful model system used to understand the mechanisms of DNAJ proteins, owing to their high conservation. The G/F domain of DNAJB6 is homologous to the specific yeast DNAJ protein, Sis1 [15]. This is convenient because, unlike DNAJB6, there are known substrates for Sis1, and the deletion of Sis1 results in strong phenotypes. For example, Sis1 is required for cell viability and is essential for yeast prion propagation [15]. Thus, studying the impact of human LGMD1D mutations in the yeast Sis1 model allows one to form hypotheses about how they might impact DNAJB6 function. Indeed, we previously expressed a chimeric Sis1-DNAJB6, in which Sis1’s G/F domain was replaced with that of DNAJB6 [15]. The yeast expressing the Sis1-G/F chimera were viable and supported prion propagation, consistent with the homology between these proteins. However, when an LGMD1D missense mutation within the G/F domain was introduced, prion propagation was impaired, suggesting a partial loss-of-function [15]. This effect on prion propagation appears to be conserved, because DNAJB6b also facilitates the disaggregation of proteins with prion-like, low complexity sequences, such as huntingtin and TDP-43, in mammalian cells [15, 16]. Moreover, LGMD1D mutations in DNAJB6 impair this function [15]. Together, these observations suggested to us that the Sis1 yeast model might be used to probe the mechanism of LGMD1D mutations.

Here, we use yeast, mammalian cells, and a new mouse model, combined with chemical probes targeting chaperones, to reveal an unexpected relationship between DNAJB6 mutations and aberrant cycling of HSP70 at the Z-disc. Counter-intuitively, we find that inhibiting the interaction of DNAJB6 with HSP70, either genetically or pharmacologically, can partially restore HSP70 mobility and rescue muscle phenotypes in treated mice. These results suggest new ways of potentially treating LGMD1D. More broadly, they provide insight into the mechanisms of myopathy-associated chaperonopathies.

## Results

To model LGMD1D mutant dysfunction, we generated homologous DNAJB6 disease-associated mutations in *Sis1*. We found that yeast expressing Sis1-F106L and Sis1-F115I (corresponding to DNAJB6-F93L and F100I) had reduced viability (Figure 1A). This was similar to previous reports demonstrating that deletion of the Sis1 G/F domain dominantly affects yeast viability in the setting of endogenous Sis1 [17]. This dominant effect can be rescued by a second *cis* mutation within the C-terminus, L268P, which was shown to abrogate Ssa1/Hsp70 interaction with Sis1 [18]. Similarly, we found that combining the L268P mutation with Sis1-F106L or F115I, restored yeast viability (Figure 1A). Notably, generating a second mutation in Sis1-F115I that impairs Sis1 dimerization, ΔDD, did not rescue viability (Figure 1A). These data indicate that the gain of LGMD1D mutant toxicity occurs via the DNAJ-HSP70 interaction and not DNAJ dimerization. Importantly, we confirmed that these effects were not due to a destabilization of Sis1 (Figure 1B).

**Figure 1:**
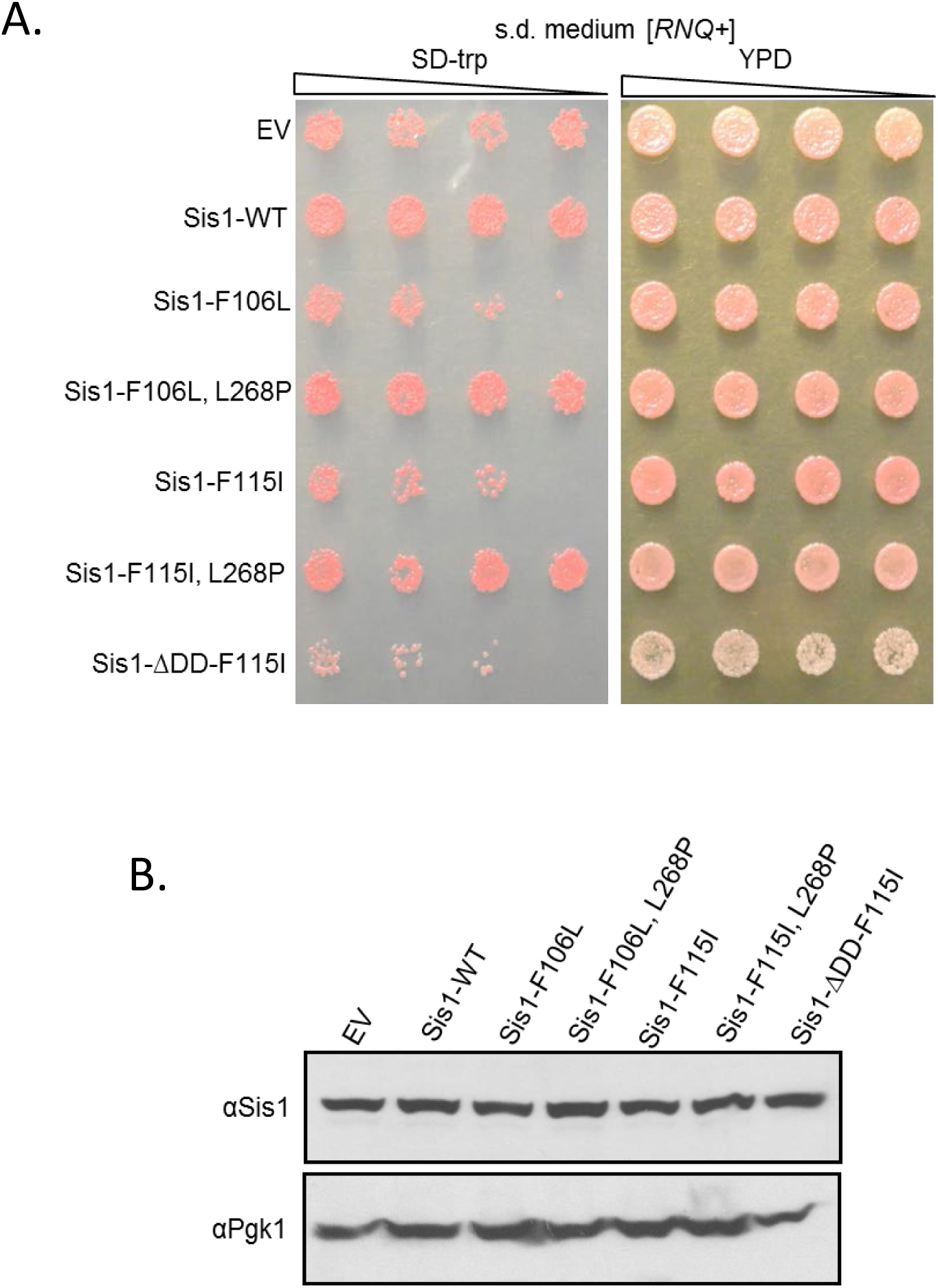
LGMD1D mutations are toxic in yeast Sis1 and rescued by impairing HSP70 association. A) Yeast cells propagating m.d. high [RNQ+] transformed with plasmids over-expressing Sis1-WT, Sis1-F106L (DNAJB6-F93L homologous), Sis1-F115I (DNAJB6-F100I homologous) constructs, an empty vector control (EV) or Sis1 mutant constructs with a second mutation that impairs HSP70 association (L268P) or blocks Sis1 dimerization (ΔDD). Cultures were normalized and serially diluted five-fold and were spotted on medium (SD-trp) to select for the plasmid or medium (YPD) that provides no selection. B) Western blot analysis from lysates of yeast expressing the Sis1 constructs in B. Pgk1 is a loading control. Images for both (A) and (B) are representative of three independent experiments.

One feature of LGMD1D muscle pathology is the formation of inclusions composed of RNA binding proteins, such as TDP-43 and hnRNAPA2B1 [4, 11]. We have previously demonstrated that expression of LGMD1D mutant DNAJB6b increases the abundance of TDP-43 positive nuclear stress granules following heat shock [15]. In light of the results with our LGMD1D yeast model, we hypothesized that this gain-of-function effect was dependent on HSP70 interaction. To test this, we expressed mCherry-tagged TDP-43 with plasmid constructs containing DNAJB6b-WT or LGMD1D mutant DNAJB6b with or without an additional H31Q mutation that abrogates HSP70 association in HeLa cells (Figure 2A-C). While LGMD1D mutant DNAJB6b expression increased the number of TDP-43 positive stress granules post heat shock, the addition of the H31Q mutation resulted in TDP-43 granules that appeared similar to DNAJB6b-WT expression (Figure 2A-B). We further tested whether the H31Q mutation ameliorated TDP-43 insolubility, as TDP-43 stress granules become detergent insoluble in the presence of LGMD1D mutant DNAJB6b [15]. In order evaluate TDP-43 aggregation, we subjected lysates to differential extraction and centrifugation generating a total lysate fraction (T), a soluble supernatant fraction (S) and an insoluble pelleted fraction (P). Similar to what was observed with fluorescence microscopy, there was an increase in insoluble/pelleted (P) TDP-43 following heat shock in cells expressing LGMD1D mutant DNAJB6b as compared to DNAJB6b-WT expression. However, when an H31Q mutation was present with the disease mutation, the increase in pelleted TDP-43 was reduced (Figure 2C). In combination with our yeast model, these data provide additional support that the DNAJB6-HSP70 interaction plays a critical role in LGMD1D pathogenesis.

**Figure 2:**
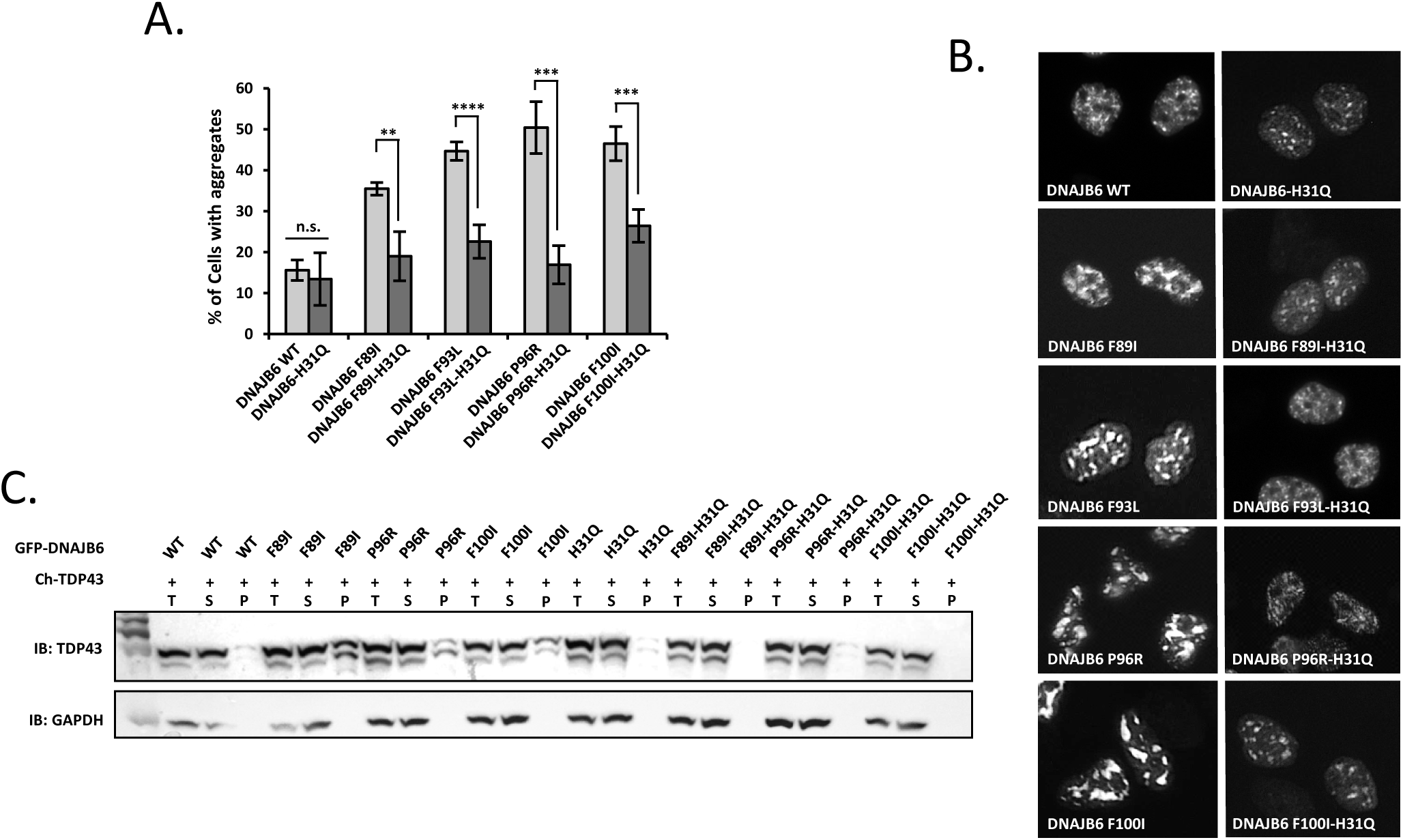
LGMD1D mutations in DNAJB6b impair TDP-43 disaggregation which is corrected with a second H31Q mutation. A-C) Hela cells were co-transfected with mCherry-TDP-43 and wild-type or LGMD1D mutant (F89I, F93L, P96R or F100I) GFP-DNAJB6b. In some cases a second mutation in the J domain (H31Q) was also present. Transfected cells were heat shocked at 42° for one hour prior to analysis. Quantification of the percentage of transfected cells with TDP-43 aggregates. The comparison between groups was performed using a paired Student t-test **P < 0.005, ***P < 0.0005, ****P < 0.00005. B) Representative fluorescent images of mCherry-TDP-43 positive nuclei quantified in (A). C) Lysates from cells in (A) were separated into total (T), soluble (S), and pellet (P) fractions and then subjected to SDS-PAGE and Western blot using an α-TDP43 and α-GAPDH antibodies.

To dissect how the H31Q mutation modulates DNAJB6 function, we specifically examined the binding of DNAJB6 and HSP70. As expected based on previous results [19], the H31Q mutation decreased the binding of DNAJB6b-WT to co-expressed Flag-HSP70 (Figure 3A). Notably, LGMD1D mutant DNAJB6b had increased association with Flag-HSP70 and this interaction was weakened by a secondary H31Q mutation (Figure 3A). Abrogating the HSP70 interaction in LGMD1D mutant DNAJB6b with the H31Q mutation also corrected other gain-of-function effects, such as an increase in LGMD1D mutant DNAJB6b stability (Figure 3B-C), and heat shock associated cell death (Figure 3D). These data suggest that the gain-of-function effect of LGMD1D mutant DNAJB6b can be partially corrected by inhibiting the DNAJB6:HSP70 interaction.

**Figure 3:**
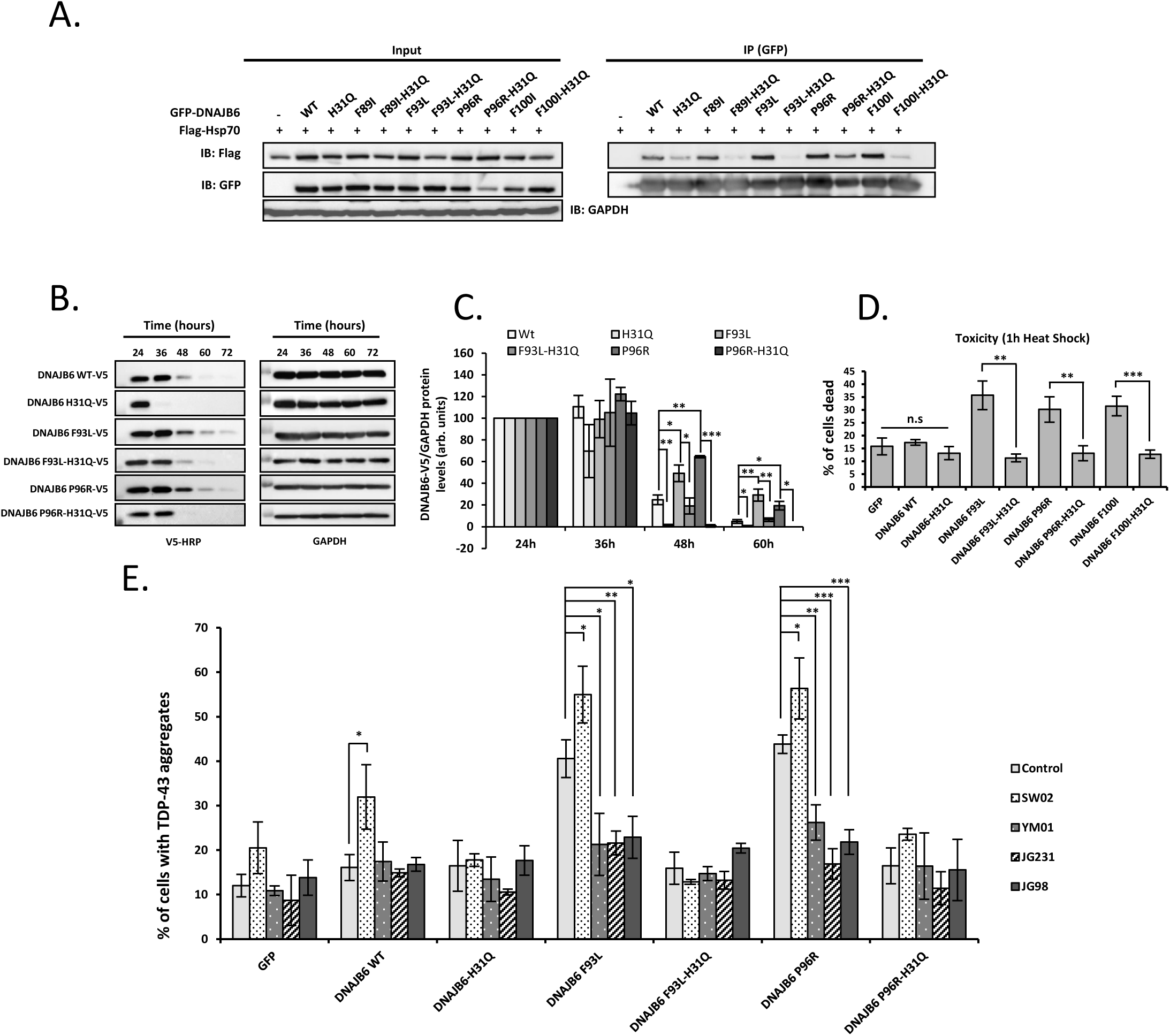
LGMD1D mutant dysfunction is corrected by abrogating HSP70 interaction. A) Hela cells were co-transfected with plasmids expressing Flag-Hsp70 and wild-type or LGMD1D mutant (F89I, F93L, P96R or F100I) GFP-DNAJB6b. In some cases a second mutation in the J domain (H31Q) was also present. 24 hours later, cells were lysed and GFP-DNAJB6 was immunoprecipitated and subsequently immunoblotted for GFP or Flag. B) Tetracycline inducible isogenic 293 cell lines expressing (WT, H31Q, F93L, F93L-H31Q, P96R or P96R-H31Q) V5-DNAJB6b were induced with tetracycline for 24h and then removed for the indicated times. Lysates were immunoblotted for V5 and GAPDH. C) Quantitation of V5-DNAJB6b from three independent experiments. comparison between groups was performed using a paired Student t-test. * P≤ 0.05, **P < 0.005 and ***P < 0.0005. D) Hela cells were transfected with the indicated constructs, subjected to heat shock at 42° for 1 hour and the percentage of ethidium homodimer-1-positive cells was quantitated. Data are presented as percent cells found positive/dead. Comparison between groups was performed using a paired Student t-test **P < 0.005, ***P < 0.0005. E) Hela cells were co-transfected with mCherry-TDP-43 and wild-type or LGMD1D mutant (F89I, F93L, P96R or F100I) GFP-DNAJB6b. In some cases a second mutation in the J domain (H31Q) was also present. After 12h of transfection, cells were treated for 16h with either YM-01, JG231, JG98 (Hsp70 inhibitors) or SW02 (Hsp70 activator). The percentage of cells with TDP-43 positive nuclear aggregates granules was quantified following 1h of heat shock. The comparison between groups was performed using a paired Student t-test *P < 0.05, **P < 0.005, ***P < 0.0005.

Based on these results, we wanted to further explore whether HSP70 might be a therapeutic target in LGMD1D. Specifically, we wondered whether inhibiting the association of HSP70 with DNAJB6b with small molecules might partially protect against LGMD1D phenotypes in mammalian disease models. As a proof-of-concept, we tested four different compounds known to modulate HSP70 interactions with co-chaperones [20]. YM-01, JG98 and JG231 inhibit HSP70 activity by interfering with its interactions with co-chaperones, including DNAJ proteins [21]. Conversely, SW02 activates HSP70 by enhancing its interaction with DNAJ proteins [22]. At 12h post-transfection with DNAJB6b and TDP-43, HeLa cells were treated for 16 hours with vehicle, or an HSP70 modulator prior to heat shock. By quantifying the percentage of cells with TDP-43 granules (Figure 3E), we found that SW02 increased the gain-of-function effect of LGMD1D mutant DNAJB6b, whereas YM-01, JG98 and JG231 reduced TDP-43 inclusions, specifically in LGMD1D mutant DNAJB6b expressing cells. These effects seemed to be dependent upon the HSP70 interaction, because there was no change when the HSP70 modulators were applied in the setting of LGMD1D mutant DNAJB6b harboring an additional H31Q mutation (Figure 3E).

We have previously described a mouse model that overexpresses V5-tagged DNAJB6b-F93L using an MCK promoter in skeletal muscle [12]. These mice had premature death, weakness, myopathic muscle and protein inclusions similar to that seen in LGMD1D patients, and displayed these phenotypes as early as two months of age [12]. To generate a model of LGMD1D that did not rely on overexpression, we generated the DNAJB6-F90I mutation (homologous to the F89I mutation in humans) in mice using CRISPR/Cas9. Heterozygous DNAJB6-F90I mice developed progressive weakness, muscle atrophy and scattered fibers with myofibrillar disorganization after 12 months of age (Figure 4A-F). Consistent with the phenotypes seen in the transgenic mouse model, immunoblot analysis showed an increase in both DNAJB6a and DNAJB6b isoforms, as well as desmin (Figure 4G). In addition, several chaperone proteins were increased, as assessed by qPCR (Figure 4H), suggesting a stress response and impairment of protein quality control. For example, the levels of both HSPA1A (HSP70) and HSPA8 (Hsc70) were elevated, as demonstrated via immunoblot and qPCR (Figure 4G-H). Notably, although DNAJB6 protein accumulated, its mRNA levels were unchanged (Figure 4H). Thus, much like what was previously observed in our transgenic models, LGMD1D mutations in DNAJB6 seem to lead to an age-associated dysfunction in protein quality control and an aberrant accumulation of Z-disc proteins [12].

**Figure 4:**
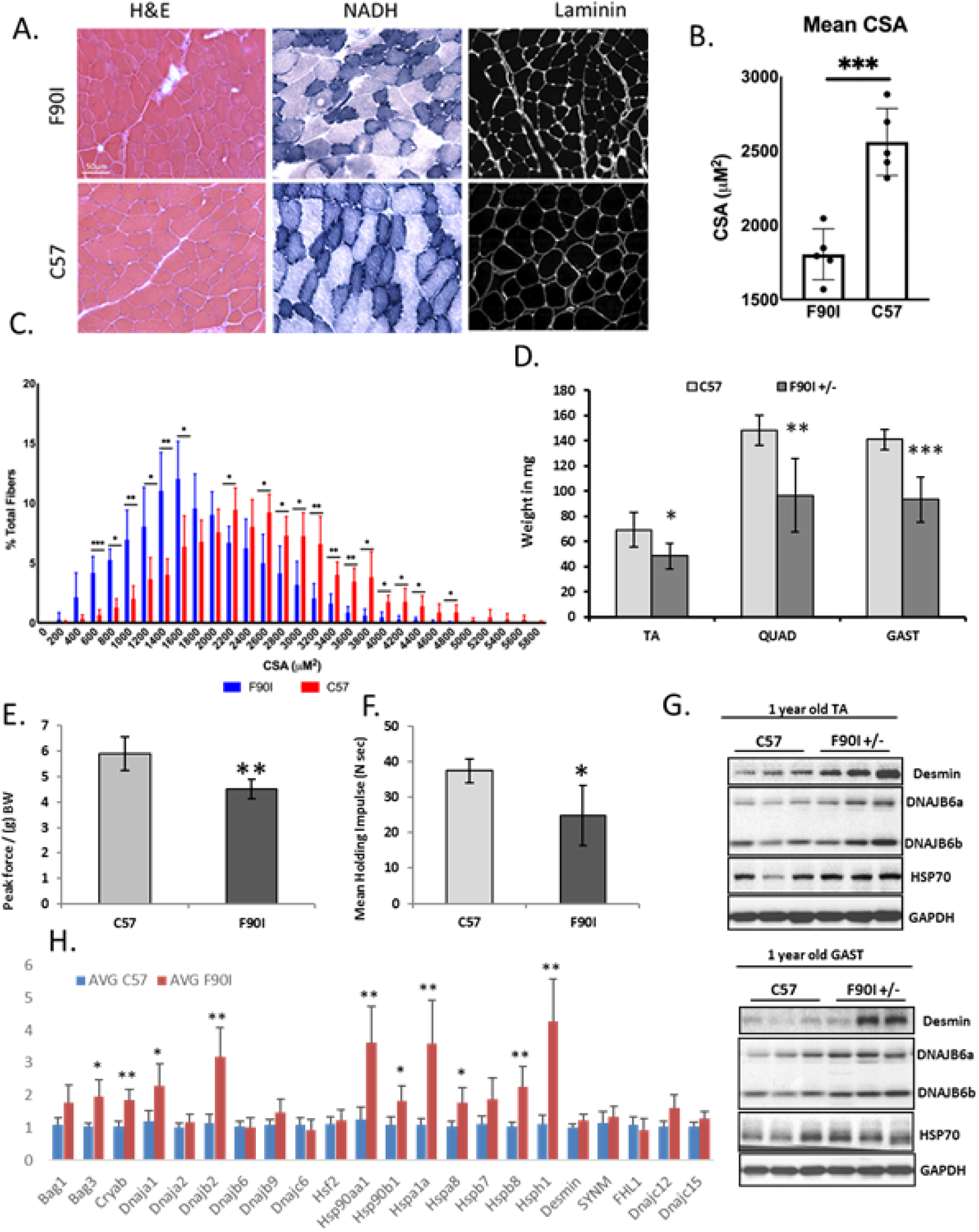
DNAJB6 F90I knockin mice develop myopathy. A) Histochemical analysis with H&E and NADH staining of tibialis anterior (TA) muscle from one year old F90I (heterozygous) or control C57 mice. Immunohistochemical analysis with an anti-laminin antibody of the same muscle. B-C) Quantitation of the cross sectional area muscle from (A). D) Graph representing the average mass (mean ± SD, weight in milligrams) of TA, Quadriceps (QUAD) and Gastrocnemius (GAST) muscles from 1 year old C57 and DNAJB6 F90I +/-mice. Comparison between groups was completed by using a paired Student t-test. *P < 0.05, **P < 0.005, ***P < 0.0005. E) Forelimb grip strength measurements from one year old littermate control (C57) or DNAJB6 F90I heterozygous mice and (F) inverted wire screen holding on the same animals in (E). Comparison between groups was completed by using a paired Student t-test. *P < 0.05, **P < 0.005. G) Immunoblot of lysates from TA and GAST muscle from three different one year old C57 and F90I heterozygous mice using antibodies to DNAJB6, Hsp70, Desmin and GAPDH. H) qPCR analysis of TA muscle from five 1 year old F90I heterozygote and C57 mice. Data are presented as a fold change as compared with C57 controls. Comparison between groups was completed by using a paired Student t-test. *p<0.05, **P < 0.005.

Because DNAJB6 normally collaborates with HSP70 to reversibly cycle onto “client” Z-disc proteins [14] and myofibrillar disorganization is a pathologic feature of LGMD1D, we hypothesized that this process might be damaged by LGMD1D mutations. To test this idea, we evaluated the localization of DNAJB6b and HSP70 in live mouse muscle by imaging the flexor digitorum brevis (FDB) following electroporation of GFP-tagged DNAJB6b or HSP70. As expected, DNAJB6b-WT localized to the Z-disc, while HSP70 localized to the Z-disc and, to varying degrees, the M-line (Figure S1A). Localizations were confirmed via second harmonic generation imaging, which identifies the A bands and by expression of desmin-GFP with mCherry-DNAJB6b-WT (Figure S1A-B). To understand chaperone dynamics at the Z-disc, we performed fluorescence recovery after photobleaching (FRAP) of DNAJB6b and HSP70. Fluorescence from DNAJB6b-WT was rapidly recovered at the Z-disc (Figure 5A-B), indicating that most of the protein is mobile (Figure 5C) and transiently associates with Z-disc clients. By contrast, two different LGMD1D mutations (F89I and P96R) in DNAJB6b showed slower recovery at the Z-disc (Figure 5A-B).

**Figure 5:**
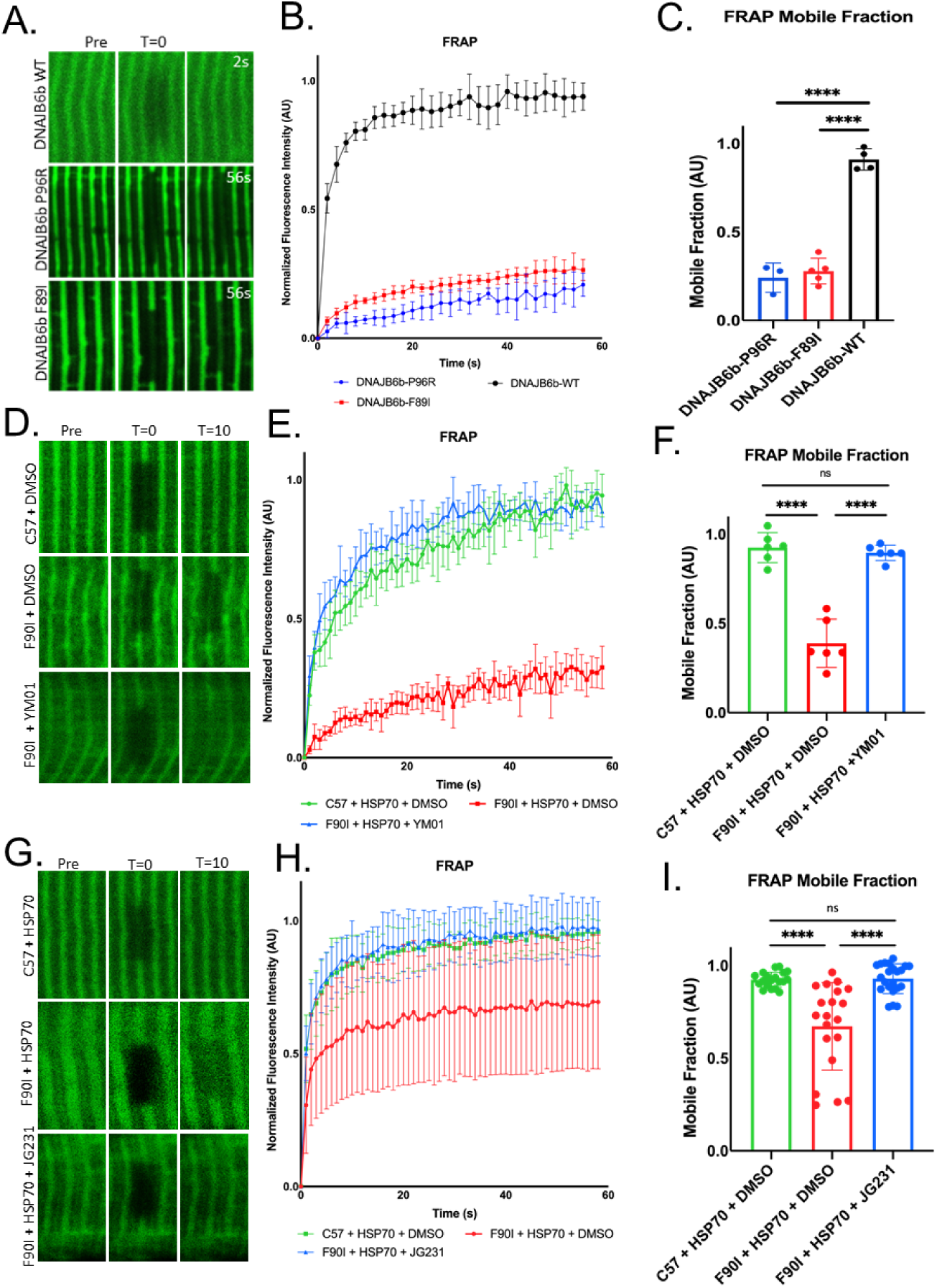
Imaging of DNAJB6 and HSP70 kinetics at the Z-disc. A) Fluorescence recovery after photobleaching was performed with two-photon microscopy on mouse foot pad following electroporation into the FDB with constructs expressing GFP-DNAJB6b-WT, -P96R, or -F89I. Representative images show baseline prior to bleaching (pre), immediately post-bleach (t=0s), and following (2s or 56s) of recovery (t=2s or t=56s). B) Graph of the normalized RFI vs time in seconds for the studies in (A). D) Graph of the percentage of maximum fluorescence recovery corresponding to the mobile vs immobile fraction. D-I) Experiments similar to (A) following electroporation of HSP70-GFP in C57 control or DNAJB6-F90I heterozygous mice. In some cases, mouse footpad was injected with YM01 (D-F) or mice were given i.p. injections of JG231 (G-I). D and G) Representative images show baseline prior to bleaching (pre), immediately post-bleach (t=0s), and following 10s of recovery (t=10s). E and H) Graph of the normalized RFI vs time in seconds for the studies in (D and G respectively). F and I) Graph of the percentage of maximum fluorescence recovery corresponding to the mobile vs immobile fraction for the studies in (D and G respectively).

Using this system, we next wanted to test whether the slow recovery of DNAJB6b-F89I might also affect HSP70 in a dominant manner. To test this concept, we performed a similar FRAP experiment using 3 month old DNAJB6-F90I knockin mice that express one copy of the DNAJB6-F90I (F89I in humans) at endogenous levels. Three months was chosen as the time point because, at this early age, the mice do not show any obvious muscle pathology. Whereas HSP70 had rapid fluorescence recovery in littermate control mice, its recovery was slowed in DNAJB6-F90I knockin mice (Figure 5D-F). Local delivery of YM-01 to the footpad (Figure 5D-F), or systemic treatment with JG231 (Figure 5G-I), increased HSP70 recovery kinetics in DNAJB6-F90I mouse muscle. Importantly, JG231 did not significantly affect the recovery kinetics of a mutant HSP70, Y149W, in which the compound-binding site is occluded, supporting the idea that the compound’s effect was mediated by binding to HSP70 (Figure S2A-C).

To explore whether abrogating DNAJB6-HSP70 interactions with JG231 might also restore muscle functions, we treated transgenic DNAJB6b-WT or DNAJB6b-F93L overexpressing mice with vehicle (DMSO) or 4 mg/kg i.p. JG231 every other day for eight weeks and measured mouse performance on forelimb grip and hanging grid tests (Figure 6A-B). Mice were sacrificed at 4 and 8 weeks for muscle histopathology (Figure 6C-D). This dose and schedule were chosen based on previous safety and pharmacokinetics studies [23]. Although DNAJB6b-F93L mice are significantly weaker than DNAJB6b-WT mice, JG231 treatment increased their performance as early as one week (Figure 6A-B), and at 4 and 8 weeks, these mice also had an increase in isolated muscle mass (Figure 7A-B). To further assess the effect of JG231 treatment, we quantitated myofiber size and degree of myofibrillar disarray in DNAJB6b-WT or DNAJB6b-F93L mice treated with or without JG231 following treatment at 4 and 8 weeks (Figure 7C-E). Myofibrillar disarray was quantitated using an NADH stain and scored from 0-6 in a blinded fashion by two neuromuscular physicians with muscle pathology expertise (AF and CW). Examples of scored images are in Figure S3. Notably, there was a significant shift toward larger myofibers and a decrease in the percentage of fibers with abnormal internal architecture, as demonstrated on NADH staining at both 4 and 8 weeks (Figure 7E).

**Figure 6:**
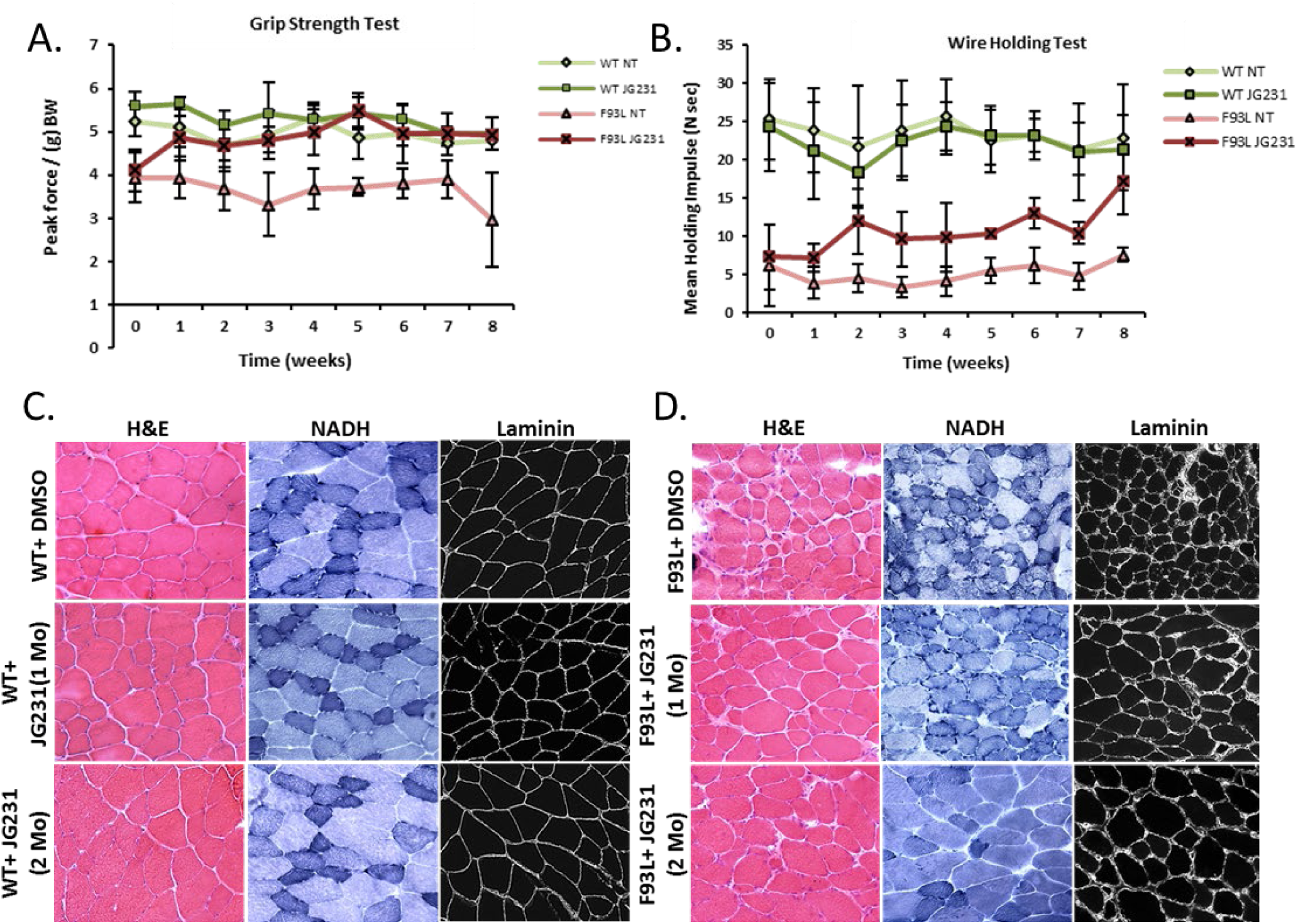
JG231 improves strength and myopathology in LGMD1D mice. Age and sex matched 4 month old DNAJB6b-WT and DNAJB6b-F93L mice were treated for 4 and 8 weeks with intraperitoneal vehicle or JG231. A) Forelimb grip strength and B) inverted wire screen holding test were performed weekly. C-D) Histochemical analysis with H&E and NADH staining or immunohistochemical analysis with anti-laminin of TA muscle from DNAJB6b-WT or DNAJB6b-F93L after 4 or 8 weeks treatment.

**Figure 7:**
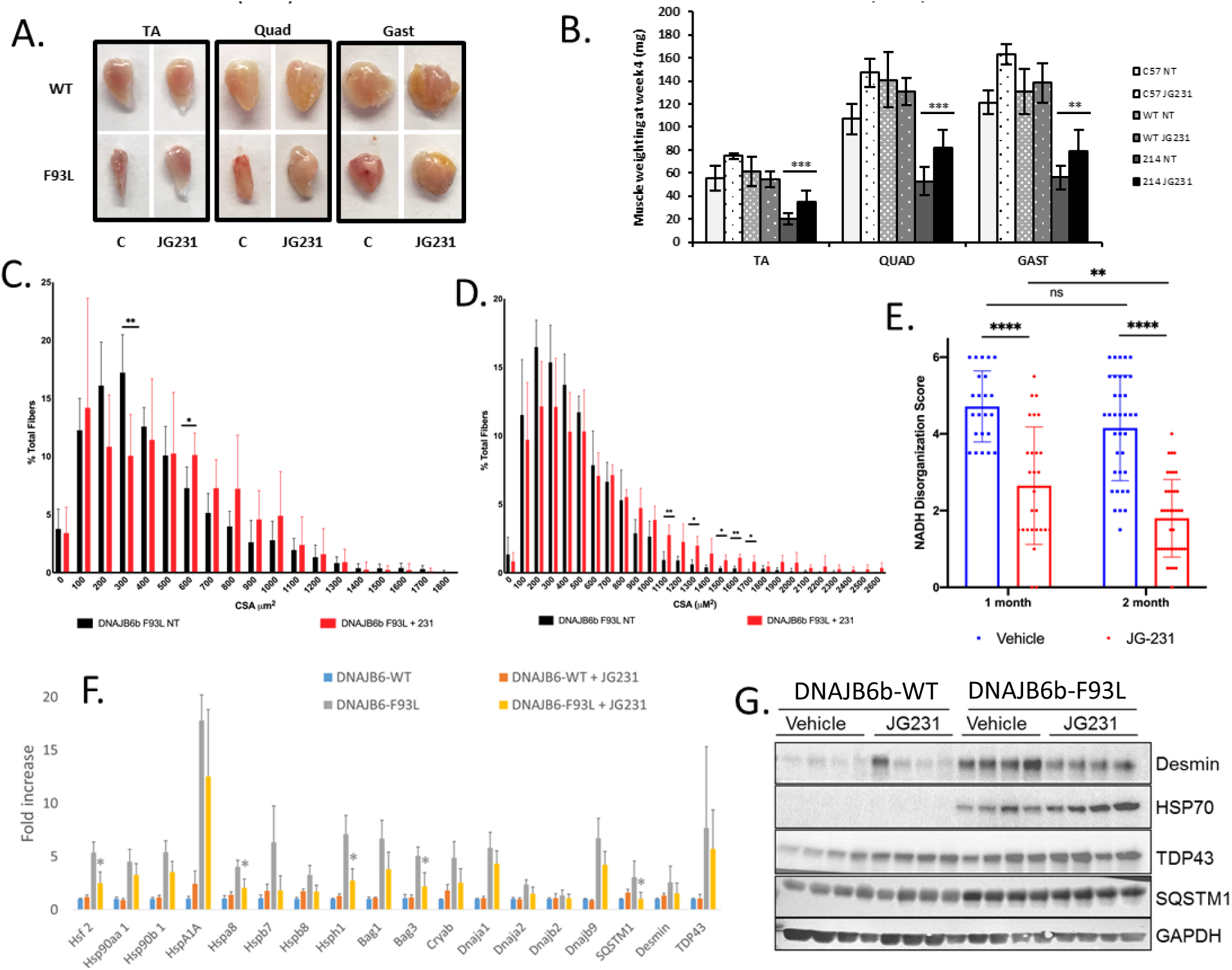
JG231 increases myofiber size in LGMD1D mice. A) Representative mages of TA, QUAD and GAST muscles from 4 month old DNAJB6b-WT and DNAJB6b-F93L mice treated with vehicle (c) or JG231 for 4 weeks. B) Graph of the average mass (mean ± SD, weight in milligrams) of the TA, QUAD and GAST from six DNAJB6b-WT and DNAJB6b-F93L mice treated with vehicle or JG231 for 4 or 8 weeks. **P < 0.005, ***P < 0.0005. We have used frozen sections of TA muscle from 4 month old DNAJB6b WT and DNAJB6b F93L mice control or treaded with JG231 for 4 and 8 weeks. C-D) Graph of the distribution of myofiber diameter in TA muscle of 4 month old DNAJB6b-WT and DNAJB6b-F93L mice treated with vehicle or JG231 for 4 weeks (C) or 8 weeks (D). E) Graph of NADH disorganization scored from TA muscle from 4 month old DNAJB6b-WT and DNAJB6b-F93L mice treated with vehicle or JG231 for 4 and 8. Error bars represent the standard error. F) qPCR analysis of TA muscle from 4 months old DNAJB6b-WT and DNAJB6b F93L treated with vehicle or JG231 for 4 weeks. Data are presented as a fold change as compared with C57 controls. Comparison between groups was completed by using a paired Student t-test. *p<0.05, **P < 0.005. G) Immunoblot of TA muscle lysates from four different DNAJB6b-WT and DNAJB6b-F93L mice treated with vehicle or JG231 for 8 weeks using antibodies to Desmin, HSP70, TDP43, SQSTM1 and GAPDH.

JG231 treatment had a modest effect on the expression of other chaperone proteins, as evaluated by qPCR of tibialis anterior muscle (Figure 7F), suggesting that this effect was independent of a global stress response. To determine if there were changes in other proteins found to accumulate in LGMD1D muscle, we immunoblotted for TDP-43, SQSTM1, HSP70 and desmin (Figure 7G). Strikingly, after 8 weeks of treatment with JG231, there was a significant decrease in total desmin protein levels (Figure 7G), as might be expected for a partial recovery of quality control. The observed decrease in desmin accumulation also matches the improvement in internal, muscle architecture that were seen in the treated mice. Together, these results suggest that the DNAJB6-HSP70 interaction is an unexpected drug target for LGMD1D.

## Discussion

Protein chaperones are important for the maintenance of sarcomeric structure and muscle function [7]. Dominant mutations in DNAJB6 cause LGMD1D, a myopathy with myofibrillar disorganization and protein inclusions [3, 4]. Our data supports an unexpected mechanism in which the dominant effect of mutations in DNAJB6’s G/F domain require its interaction of HSP70. Specifically, yeast expressing homologous LGMD1D mutations in Sis1 have reduced viability, but this toxicity is corrected by including a second mutation that abrogates binding to HSP70. Moreover, genetically blocking the interaction DNAJB6 with HSP70 rescues the dominant effect of LGMD1D mutations on TDP-43 disaggregation and post-heat shock toxicity. There are a number of possible mechanisms to link these observations. For example, the interactions of mutant DNAJB6 with HSP70 might reduce the stability of DNAJB6 clients. HSP70 has been shown to prevent proper folding if it is not allowed to cycle off client [24]. Indeed, although HSP70 is often associated with its roles in protein folding, its prolonged binding appears to be antagonistic to this process in many settings {Sekhar, 2012 #305}. Alternatively, the prolonged interaction of mutant DNAJBb with HSP70 might titrate it away from other clients, leading to broader disruption of HSP70-dependent processes [25]. These mechanisms are not mutually exclusive, and both could result in a more general impairment of protein quality control.

Several lines of evidence suggest that Z-disc proteins, such as desmin, may be DNAJB6b clients. For example, loss of DNAJB6 in myoblasts leads to the accumulation of desmin [14]. We found that normal DNAJB6b and HSP70 transiently localize to the Z-disc in mouse muscle, as measured by FRAP. However, in mice expressing an LGMD1D mutation, the movement of HSP70 on the Z disc is slowed, suggesting that it is more avidly associated with mutant DNAJB6. Strikingly, treatment with small molecules known to interrupt DNAJ and HSP70 interactions [21] partially recovered HSP70 kinetics. These results further substantiate the idea that the phenotypes associated with LGMD1D mutant DNAJB6 require HSP70 interactions.

Moreover, a recent study exploring the pathogenic mechanism of mutations in BAG3 associated with another chaperonopathy showed striking similarities [26]. Dominant mutations in BAG3 lead to its destabilization and subsequent aggregation [27]. Mutant BAG3 aggregates contain additional chaperones, including HSP70 and DNAJB6. Likewise, blocking the interaction of HSP70 with BAG3 can partially rescue phenotypes, suggesting that BAG3-associated myopathy is, like LGMD1D, due to a dominant interaction with HSP70. The similarities between these findings suggest that myopathies might share a common molecular feature of aberrant protein quality control, leading to a “traffic jam” of chaperones on Z-disc proteins. Re-mobilizing these chaperones might be a generalizable approach to partially restore muscle function in myopathies.

More immediately, this study suggests that the HSP70–DNAJB6 interaction could be a drug target for the treatment of LGMD1D. However, while HSP70 inhibitors have been studied in preclinical safety studies in mice [28, 29], any treatment for human myopathy would require long-term administration and the safety of chronic treatment is not yet clear. The key to translation of this concept might be in creating more selective inhibitors, that act exclusively on the DNAJB6–HSP70 interaction. The inhibitors currently available, while useful chemical probes for proof-of-concept studies, have broad effects on HSP70 co-chaperone contacts. Thus, a selective inhibitor of mutant DNAJB6–HSP70 interactions may be the (safest and) most effective.

## Methods

### Yeast Studies

The yeast strains used in this study are derived from *Saccharomyces cerevisiae* 74-D694 (*ade1-14 his3-Δ200 leu2-3, 112 trp1-289 ura 3-52*). Yeast grown and manipulated using standard techniques. As indicated, cells were grown in rich media YPD (1% yeast extract, 2% peptone, 2% dextrose) or in synthetic defined (SD) media (0.67% yeast nitrogen base without amino acids, 2% dextrose) lacking specific nutrients to select for appropriate plasmids. Wild-type (WT) yeast harboring the s.d. medium [*RNQ*+] variant and the [*rnq*-] control strain were kindly provided by Dr. S. Liebman [30]. Construction of *ΔSis1* [*rnq-*] and s.d. medium [*RNQ+*] yeast strains were described previously [15]. Medium containing 1mg/mL 5-fluoroorotic acid (5-FOA) that select against cells maintaining *URA3*-marked plasmids was used to replace WT Sis1 with the mutant constructs using the plasmid shuffle technique. Plasmid transformations were done using PEG/LioAC technique and the cells were selected using SD-trp plates. Plasmids pRS316-*SIS1* and pRS314-*sis1-L268P* were received as gifts from Dr. E. Craig [17]. Construction of pRS314-*SIS1* was described earlier [15]. Using pRS314-*SIS1* as template, mutations in *SIS1* were generated using bridge PCR with the indicated oligonucleotides: *sis1*-*F106L* (5’GATTCTCCGGAGGACATGCGCTCAGTAATGAGGATGC and 5’GCATCCTCATTACTGAGCGCATGTCCTCCGGAGAATC), *sis1-F115I* (5’GATGCTTTCAATATTATTTCACAATTCTTTGGC and 5’GCCAAAGAATTGTGAAATAATATTGAAAGCATC). Using pRS314-*sis1-F106L* and pRS314-*sis1-F115I* as templates, double mutations with *sis1-L268P* were generated Using (5’GTTTCTCTAGTTATCCATCTG and 5’GTTTCTCTAGTTATCCATCTG) primer pair. pRS414*GPD-sis1-ΔDD-F115I* was created by using (5’ATACTAGTATGGTCAAGGAGACAAAAC and 5’CGCATCGATTTATGGATAGTCCACTTTATATTTTAC) primers, followed by digestion with SpeI/ClaI, and ligation with pRS314*GPD* that was digested with the same enzymes. Yeast cells were spotted as described previously [31]. Briefly, yeast cells grown overnight were pelleted, washed and normalized with water to an optical density of 1.0. The cells were serially diluted (1:5) using a multichannel pipette into a 96-well plate and were spotted onto agar plates using an ethanol-sterilized 48-pin replicator. For protein analysis, yeast cells were lysed by vortexing with glass beads in buffer containing 100mM Tris-HCl pH 7.5, 200 mM NaCl, 1 mM ethylendiaminetetraacetic acid (EDTA), 5% glycerol, 0.5 mM dithiothreitol (DTT), 50 mM N-ethylmalemide (NEM), 3 mM phenylmethanesulphonylfluoride (PMSF) and complete Mini protease inhibitor cocktail (Roche). Following lysis, an equal volume of RIPA buffer (50 mM Tris-HCl pH 7, 200 mM NaCl, 1% Triton X-100, 0.5% sodium deoxycholate, 0.1% sodium dodecyl sulfate (SDS) was added to the lysate and centrifuged briefly to obtain the total protein fraction. Protein concentrations were normalized and subjected to SDS-PAGE, transferred to PVDF membranes and probed with an anti-Rnq1p antibody. The total protein fraction was also probed with anti-Sis1p and anti-pgk1 antibodies.

### qPCR

Total RNA was isolated from TA muscle with SV Total RNA isolation kit (Promega, Z3100) according to the manufacturer’s instructions. The concentration and quality of the total RNA isolated was determined using a Nanodrop spectrophotometer. cDNA was synthesized using the Transcriptor first strand cDNA synthesis kit (Roche, 04379012001). Gene expression levels were analyzed by real-time PCR on an Applied Biosystems model 7500 (software v2.0.5) using the Faststart universal SYBR Green master ROX qPCR mastermix (Roche, 04913850001). Quantitative polymerase chain reaction was performed with primers to Chaperone proteins (CRYAB, Hsp and BAG families), Cytoskeletal proteins (Desmin, SYNM), FHL1, HSF, TDP43, SQSTM1 and hDNAJA/B/C families. The values were normalized to GAPDH and represented as fold change. Primer sequences are in Supplemental Material.

### Antibodies

Antibodies used were the following: anti-rabbit Sis1 (Cosmo Bio Co., ltd., COP-080051), anti-mouse Pgk1 (Abcam, 113687), anti-rabbit GAPDH (Cell Signaling, 2118), anti-V5 HRP Invitrogen (B96125), anti-mouse Desmin (Dako, M0760), anti-rabbit TDP43 (Proteintech, 10782-2-AP), anti-rabbit DNAJB6 (abcam, ab198995), anti-mouse Hsp70 (Enzo, ADI-SPA-812), anti-rabbit P62 “SQSTM1” (Proteintech, 18420-1-AP), anti-rabbit laminin (ab1575), anti-rabbit GFP (Sigma g1544), anti-mouse Flag (Sigma F3165). Secondary antibodies include anti-mouse HRP (Pierce), anti-rabbit HRP (cell signaling) anti-rabbit AlexaFluro (555 and 488).

### Plasmids and constructs

Mammalian constructs of DNAJB6b were cloned using site-directed mutagenesis, digested with HindIII/XhoI, and ligated into vector pcDNA3.1 containing a green fluorescent protein (GFP) tag. DNAJB6b H31Q, F93L, P96R, F100I, F93L-H31Q, P96R-H31Q and F100I-H31Q mutations were generated with the Quick Change Mutagenesis Kit (Agilent Technologies #200517). Human TDP-43 fused to pCherry was described previously [32]. GFP-Hsp70 plasmid was obtained from addgene (15215).

### Western blotting

Muscle tissues and cultured cells were homogenized using RIPA lysis buffer (50 mM Tris–HCl, pH 7.4, 150 mM NaCl, 1% NP-40, 0.25% Na-deoxycholate and 1 mM EDTA) supplemented with protease inhibitor cocktail (Sigma-Aldrich), and lysates were centrifuged at 13 000 rpm for 10 min. Protein concentrations were determined using a BCA protein assay kit (Thermo Fisher Scientific). Aliquots of lysates were solubilized in Laemmli sample buffer and equal amounts of proteins were separated on 12% SDS-PAGE gels. Proteins were transferred to nitrocellulose membrane and then blocked with 5% nonfat dry milk in PBS with 0.1% Tween-20 for 1 h. The membrane was then incubated with primary antibodies, specific to the protein of interest, in 5% nonfat dry milk in PBS with 0.1% Tween overnight at 4°C. After incubation with the appropriate secondary antibody conjugated with horseradish peroxidase, enhanced chemiluminescence (GH Healthcare, UK) was used for protein detection. Immunoblots were obtained using the G:Box Chemi XT4, Genesys Version 1.1.2.0 (Syngene). Densitometry was measured with ImageJ software (National Institute of Health).

### Solubility assay

Hela cells were collected and homogenized in 250 μl of 2% SDS-radioimmunoprecipitation assay buffer (50 mg Tris–Cl (pH 8.0), 150 mg NaCl, 1% NP-40, 0.5% sodium deoxycholate and 2% SDS) and protease inhibitor cocktail (Sigma-Aldrich). Homogenates were precleared with a 30s low speed spin and an aliquot of the supernatant was collected and named the total fraction. The additional supernatant was centrifuged at 100 000 × g for 30 min at 4°C and this supernatant was collected and named the soluble fraction. The pellet was then sonicated on ice after the addition of 150 μl of 5 M guanidine-HCl and re-centrifuged at 100 000 × g for 30 min at 4°C. This supernatant was removed and named the insoluble fraction. The insoluble fraction was precipitated by adding an equal volume of 20% trichloroacetic acid (Sigma-Aldrich) and incubated on ice for 20 min. Samples were then centrifuged at 10 000 × g for 15 min at 4°C and the resulting pellet was washed twice with ice-cold acetone. Residual acetone was removed by drying tubes at 95°C and the samples were resuspended in 50 μl of 5% SDS in 0.1N NaOH. The protein concentrations of all samples were determined using a BCA protein assay kit (Pierce). Each sample (30 μg) was analyzed by Western blotting for each fraction.

### Immunoprecipitation

For immunoprecipitation of GFP-tagged DNAJB6, Hela cells were co-transfected with FLAG-Hsp70 and different DNAJB6 plasmids for 24 h. Cell pellets were collected, washed twice with cold PBS, lysed in co-IP buffer (1× tris buffered saline, 1 mM NaCl, 1% Triton X-100, 10% glycerol supplemented with phosphatase and protease inhibitors) for 15 min, and spun at 12 000g for 25 min. Lysate was collected and protein concentration was determined using Micro BCA protein assay kit. 1000 mg of total proteins from whole cell extracts were immunoprecipitated overnight at 4°C using GFP-conjugated magnetic beads (Sigma-Aldrich), then beads were washed three times with immunoprecipitation (IP) wash buffer (20 mM Tris, pH 7.5, 1 mM ethylenediaminetetraacetic acid, 150 mM NaCl, 10% glycerol). GFP-conjugated magnetic beads were resuspended in 40 ml of 2× SDS loading buffer with b-mercaptoethanol and heated at 90°C for 5 min to elute the samples. Equal amounts of total proteins (35 mg/sample) and 20 ml of eluted IP samples were resolved in 10% SDS–PAGE and analyzed in western blotting according to standard procedures.

### Wire screen holding and grip test

Grip strength testing consisted of five separate measurements using a trapeze bar attached to a force transducer that recorded peak-generated force (Stoelting, WoodDale, IL, USA). Mice have the tendency to grab the bar with their forepaws and continue to hold while being pulled backwards by the tail, releasing only when unable to maintain grip. The resulting measurement was recorded and the average of the highest three measurements was determined to give the strength score. For every time point and strain, at least five animals were used. P-values were determined by a paired Student’s t-test. To validate our results, another quantitative strength measurement was performed by wire screen holding test. Mice were placed on a grid where it stood using all four limbs. Subsequently, the grid was turned upside down 15 cm above a cage. Latency for the mouse to release the mesh is recorded and the average hanging time of three trials was used as an outcome measure.

### Animal and experimental protocols

Human DNAJB6b-WT were obtained via Addgene and the F93L point mutation was generated via QuickChange Site-directed mutagenesis by changing the cDNA position 277T>C. Both cDNAs were subcloned into a 1256MCKCAT transgenic targeting vector obtained from Dr. Stephen Hauschka, University of Washington. The promoter and coding sequence were confirmed by DNA sequence analysis. A linear fragment containing the MCK-V5-hDNAJB6 sequence was isolated by digesting the targeting vector with HindIII and KpnI and subsequent gel purification. This fragment was sent to the mouse genetics core facility at Washington University for transgenic animal production. Animals were screened for transgene insertion using PCR amplification of tail DNA. For the CRISPR/Cas9-mediated knock-in F90I mice, we used the primers. PC484.DS.Dnajb6.F - 5’ ggcaaataagactcttgggct 3’, PC484.DS.Dnajb6.R - 5’ acggaccaggcaggtactta 3’ and the sgRNA (SM721.Dnajb6.g4) TCATTGGGCAGTGTTATCATNGG. The generation of these mice was carried out by the mouse genetics core facility at Washington University for transgenic animal production. Both animal lines were bred to C57/B6 (Jackson Laboratories) to at least the F5 generation for phenotypic analysis. Control animals were non-transgenic/ F90I -/- littermates. All animal experimental protocols were approved by the Animal Studies Committee of Washington University School of Medicine. Mice were housed in a temperature-controlled environment with 12 h light–dark cycles where they received food and water ad libitum. Mice were euthanized and skeletal muscle was dissected.

### In vivo Electroporation

Mice are anesthetized using 2.5% isoflurane. We will be injecting the FDB (flexor digitalis brevis). The foot pad is wiped with ethanol and 10uL of hyaluronidase at a concentration of 2mg/mL is injected into the FDB from the base of the pad towards the digits. The mouse then is monitored in a recovery cage for 1-2hrs. Again, the animal is anesthetized with 2.5% isoflurane and the foot pad wiped with ethanol. The plasmid is then injected with endotoxin free plasmid diluted in sterile PBS to a volume of 50uL into FDB using a 29 gauge × 1/2 needle. Immediately after that, two-needle array electrodes are inserted longitudinally relative to muscle fibers. In vivo electroporation parameters for FDB are as follows: 20 pulses, 20ms duration at 1Hz, at 100 V/cm. Procedure is repeated on contralateral FDB. Needles are removed and foot is wiped again with ethanol. Mouse is then return to recovery cage for monitoring.

### Cell culture, transfection and Heat Shock

Flp-In T-REX 293 Cells (Invitrogen) expressing pcDNA5/FRT/TO constructs V5-DNAJB6b_WT, V5-DNAJB6b-F93L, V5-DNAJB6b-P96R, V5-DNAJB6b-H31Q, V5-DNAJB6b-F93L-H31Q were cultured in DMEM containing 4 mM l-glutamine (Invitrogen; 11965-084), 10% FBS (Atlanta Biologicals; S10350) and penicillin (50 IU)/streptomycin (50 μg/ml) (Invitrogen; 15 140), 50 μg/ml hygromycin B (Invitrogen; 10687-010), 50 μg/ml blasticidin (life technologies; R21001) and induced with 1 μg/ml tetracycline hydrochloride (Sigma; T76600) 48 h prior to experimentation. Cells were maintained in 5% CO2 at 37°C in 60 mm tissue culture-treated plates until the cells were 80–85% confluent. HeLa cells were cultured in high glucose formulation of Dulbecco’s modified Eagle’s medium (Sigma-Aldrich) supplemented with 10% (vol/vol) fetal bovine serum, 2 mM L-glutamine and penicillin/streptomycin. Transfection was performed with Lipofectamine 2000 (Life Technologies, Thermo Fisher Scientific; catalog 11668019) according to the manufacturer’s instructions. For heat shock experiment, HeLa cells were transfected either with GFP-DNAJB6 plasmids or co-transfected with Ch-TDP43 constructs for 24 h. After transfection, cells were subjected to heat shock (42°C, 5% CO2) for 1 h. For the drug treatment, Hela cells were treated for 16h with either DMSO, SW02 (20μM), YM01 (1μM), JG231 (10μM) or JG98 (10μM).

### Histochemistry, immunohistochemistry and microscopy

Isolated muscle was mounted using tragacanth gum (Sigma, G1128) and quick frozen in liquid nitrogen-cooled 2-methylbutane. Samples were stored at −80°C until sectioning into 10-μm sections. Hematoxylin and eosin (H&E), nicotinamide adenine dinucleotide diaphorase (NADH), and Laminin staining was performed as previously described [12]. Sections were blocked in PNB (PerkinElmer), incubated with primary antibody followed by the appropriate secondary antibody. Briefly, muscle sections were affixed to slides incubated for 10 min in ice-cold acetone, mounted with Mowiol 4–88 (sigma) + DAPI and examined using a fluorescence microscope (Nikon 80i upright+ and Roper Scientific EZ monochrome CCD camera with deconvolution software analysis [NIS Elements, Nikon]). Non-fluorescent images were taken with a 5 megapixel color CCD (Nikon). Image processing and analysis were done with (NIS Elements 4.0) software and Adobe Photoshop. Fluorescent Images of random fields were taken with a ×20 objective using a Nikon Eclipse 80i fluorescence microscope.

### Fluorescence Recovery After Photobleaching (FRAP) and Second harmonic generation

The mice were treated as follow: YM01 was diluted in DMSO to 24 mg/mL to mimic conditions used in cell culture. 10μL of YM01 or vehicle was injected into the mouse footpad 24-48 hours prior to imaging. JG2321 was diluted in DMSO to 8 mg/mL to mimic conditions used in cell culture. 50μL of JG231or vehicle was injected intraperitoneally every other day for 2 weeks prior to imaging. For imaging, Mice were anesthetized with isoflurane with 5% used for induction and 1–1.5% for maintenance during surgery at the level of reflex suppression. Skin overlying the FDB was removed. Mice were mounted on a customized viewing pane to facilitate imaging with the inverted two photon microscope. Electroporated skeletal muscle in live mice was observed with a Zeiss LSM 880 Airyscan confocal microscope with a motorized inverted Zeiss Axio Observer Z1 microscope frame. We used a 40× 1.2 NA water immersion objective, an excitation wavelength of 950 nm and an emission filter of 499-571 nm for GFP probes. Regions of interest (2.4μM × 8.49μM) were bleached with the Zeiss software. Laser settings were adjusted depending on myofiber depth to achieve at least 60% reduction in GFP signal in myofibers with variable tissue depths. This was achieved with an excitation wavelength of 950nm at 2-10% laser power for 2-5 iterations at 0.24μs duration. One image was captured immediately before and after photobleaching, followed by a series of post bleach images every 1 – 2 seconds, depending on the experiment. FRAP experiments for each condition used at least 3 different myofibers per animal. Each myofiber was tested at 3 different locations along its length. A single excitation wavelength at 800 nm was used for 2-photon electron microscopy (2PEM) and recorded using a 40× 1.2 NA water immersion objective. Second harmonic generation was detected with a non-descanned-detector through a bandpass filter of 380-430 nm. As described previously, second harmonic generation occurs mostly from 2PEM of non-centrosymetric structures, and is restricted to collagen fibers and striated muscle myosin rod domains [33]. Data were analyzed using ImageJ. We measured the average fluorescence intensity of the bleached Z-disc, a non-bleached Z-disc as a control, and a background region for each time point. The background intensity was subtracted from the bleached and unbleached intensity values. A bleached / non-bleached ratio was calculated between the corrected intensities to account for photobleaching from image capture at each time point. The value from the initial bleach image (t = 0) was then subtracted from each time point and the data were normalized so that the pre-bleach intensity was set equal to 1 and the intensity of the initial bleach image (t = 0) was set to 0. The normalized recovery data from at least 3 Z-discs per myofiber were used to generate a myofiber average. At least 3 different myofibers per animal were then averaged for further quantitative analysis. Grouped data are presented as the mean and error bars represent standard deviation. Mobile fraction was calculated from the FRAP curve as previously described [34] with some modifications. Prism 8 software was used for non-linear regression analysis. FRAP data was fit to an exponential model with one-phase association: Y=Y0 + (Plateau-Y0)*(1-exp(-K*x)). Y0 is the Y value when X (time) is zero. Plateau is the Y value at infinite time, and represents the mobile fraction. K is the rate constant. Tau is the time constant. Half-time is computed as ln(2)/K. A mobile fraction average was calculated for myofiber, and at least 3 different myofibers per animal were then averaged for further quantitative analysis. Grouped data are presented as the mean and error bars represent standard deviation. Statistical significance was assessed via unpaired t-test.

