## supplemental figures for "Inhibition of DNAJ-HSP70 interaction improves strength in muscular dystrophy"

### Figure S1

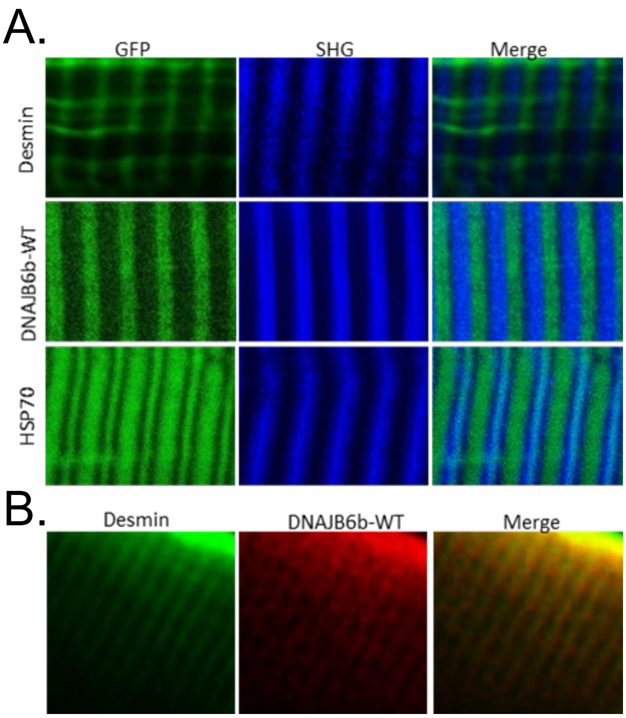


A) Desmin-GFP, GFP-DNAJB6b-WT and HSP70-GFP were electroporated in mouse FDB muscle and live mouse footpad was imaged via two-photon microscopy. Second harmonic generation was concurrently imaged to visualize the A-band for a reference point. B) Co-electroporation of Desmin-GFP and mCherry-DNAJB6b-WT mouse FDB.

### Figure S2

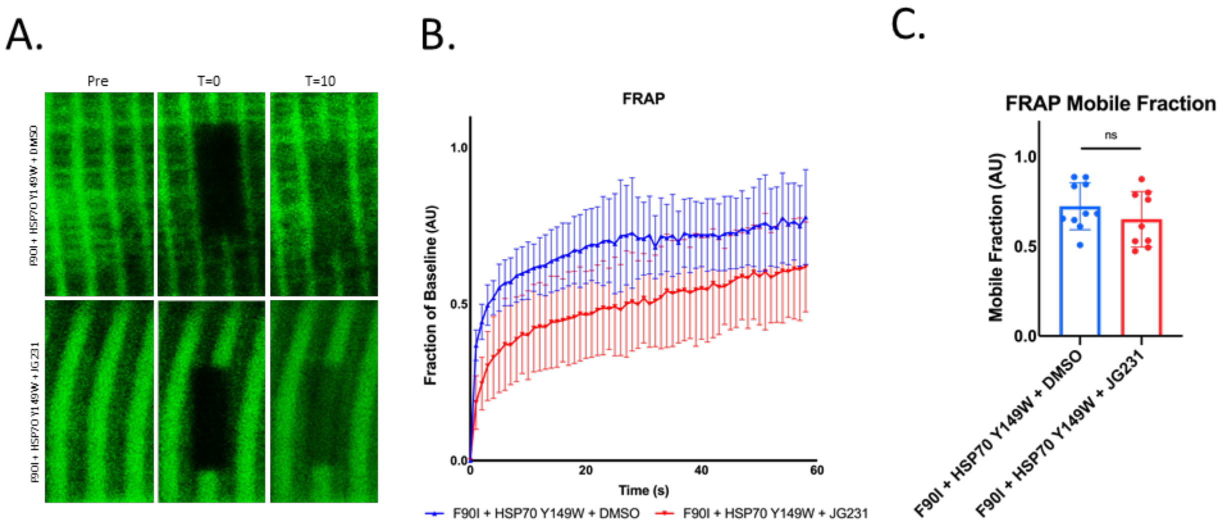


A-C) Fluorescence recovery after photobleaching was performed with two-photon microscopy on mouse foot pad following electroporation into the FDB with constructs expressing HSP70-Y149W-GFP into DNAJB6-F90I heterozygous mice. In some cases, mice were giving i.p. injections of JG231. A) Representative images show baseline prior to bleaching (pre), immediately post-bleach (t=0s), and following (10s) of recovery (t=10s). B) Graph of the normalized RFI vs time in seconds for the studies in (A). C) Graph of the percentage of maximum fluorescence recovery corresponding to the mobile vs immobile fraction from (A)

### Figure S3

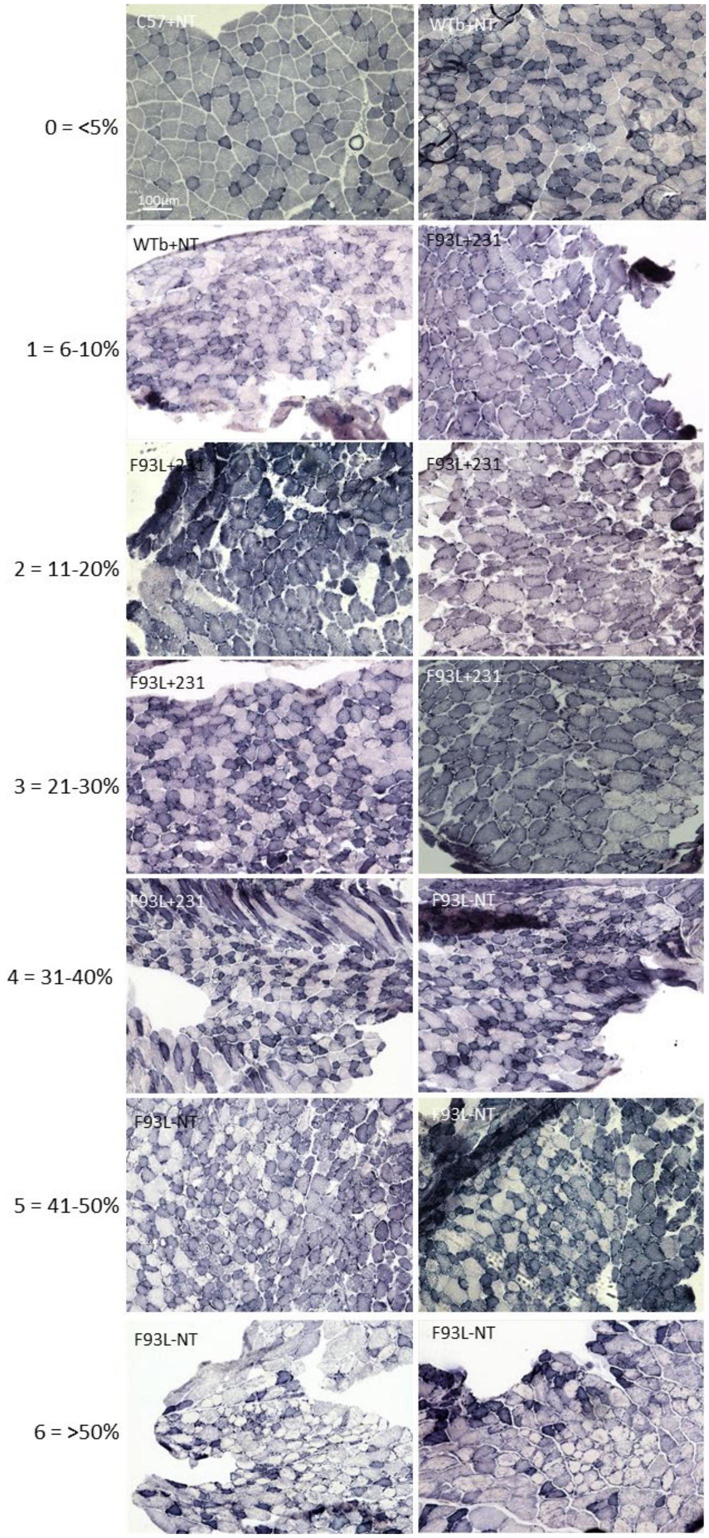


Examples of NADH myofibrillar disorganization scoring key.

### Figure S4

| | |
|---|---|
| **Hspb8 rev-qPCR mouse** | TTG GTG AAG TTC TTG GAG ACA AT |
| **Hsph1 fwd-qPCR mouse** | AAC CCC AGA TGC TGA CAA AG |
| **Hsph1 rev-qPCR mouse** | CCA CCT TTA TTT TAG GTT TCT TGG |
| **GAPDH-Left** | ATG GTG AAG GTC GGT GTG A |
| **GAPDH-Right** | AAT CTC CAC TTT GCC ACT GC |
| **Bag 1-fwd qPCRmouse** | GCT AAC CAC CTG CAA GAA TTG |
| **Bag 1-rev qPCR mouse** | TTG CAA TTC CTT AGC CAG AAA |
| **Bag 3-fwd qPCR mouse** | CCA ACT GCT CAT GGA CCT G |
| **Bag 3-rev qPCR mouse** | GCC GAG GAG GAA GAG GAT |
| **Cryab-fwd qPCR mouse** | ACG GCA AGC ACG AAG AAC |
| **Cryab-rev qPCR mouse** | TCC GGT ACT TCC TGT GGA AC |
| **Dnaja1-fwd qPCR mouse** | TGG CTC TGC AAA AGA ATG TG |
| **Dnaja1-rev qPCR mouse** | TGA ATC CTT ATC TGC ATA CCT GTC |
| **Dnaja2-fwd qPCR mouse** | TGG ATC AAC CCA GAC AAA CTT |
| **Dnaja2-rev qPCR mouse** | CTC CAA TAA CAT TAG GAA CTT CTG G |
| **Dnajb2-fwd qPCR mouse** | AGC TCG CCA TGG CTT ACA |
| **Dnajb2-rev qPCR mouse** | TGG AAC TGC AGC AAC TCT GT |
| **Dnajb9-fwd qPCR mouse** | CAC AAA GAT GCC TTT TCT ACC G |
| **Dnajb9-rev qPCR mouse** | TTA AAC TTT TCA GCT TAA TGA CGT G |
| **Dnajb6 rev-qPCR human** | TTC ATA TGC CTC CGC TAC TTG |
| **Dnajb6 fwd-qPCR human** | ATG AAG TTC TAG GCG TGC AG |
| **p62(Sqstm1)-Left** | GAA GCT GCC CTA TAC CCA CA |
| **p62(Sqstm1)-Right** | TGG GAG AGG GAC TCA ATC AG |
| **FHL1 mouse-Left** | AAGTGTGCTGGATGCAAGAA |
| **FHL1 mouse-Right** | GGGTGGCTCACTCTTGACAC |
| **DES mouse qPCR fwd** | TGCAGCCACTCTAGCTCGTA |
| **DES mouse qPCR rev** | TGAAGCTCACGGATCTCCTC |
| **SYMN mouse qPCR fwd** | AGCTCCTATCCCAGACAAGGT |
| **SYMN mouse qPCR rev** | CGACACTTTGGTGTGCTCAG |
| **TDP43 mouse qPCR fwd** | AGCATTAACCCAGCGATGAT |
| **TDP43 mouse qPCR rev** | ATGCCCATCATACCCCAAC |
| **DNAJC12 mouse fwd** | GAGGACTACTACGCCTTGCTG |
| **DNAJC12 mouse rev** | AATTCTGCCAAGATTTGCTCA |
| **DNAJC15 mouse fwd** | CCGACATCGACCACACAG |
| **DNAJC15 mouse rev** | AACAGCTGCAACACCTAGTCC |
| **Dnajc6 fwd-qPCR mouse** | GGC TCT CCG GGT GTA AAG A |
| **Dnajc6 rev-qPCR mouse** | CAT AGC TGG GCT CCA TGT CT |
| **Hsf 2 fwd-qPCR mouse** | ACC CAC ACC AAC GAG TTC AT |
| **Hsf 2 rev-qPCR mouse** | TGC TCA TCC AAG ACC AGA AA |
| **Hsp90aa 1 fwd-qPCR mouse** | GTC TCG TGC GTG TTC ATT CA |
| **Hsp90aa 1 rev-qPCR mouse** | CAT TAA CTG GGC AAT TTC TGC |
| **Hsp90b 1 fwd-qPCR mouse** | AGG GTC CTG TGG GTG TTG |
| **Hsp90b 1 rev-qPCR mouse** | CAT CAT CAG CTC TGA CGA ACC |
| **Hspa1a fwd-qPCR mouse** | GGC CAG GGC TGG ATT ACT |
| **Hspa1a rev-qPCR mouse** | GCA ACC ACC ATG CAA GAT TA |
| **Hspb8 fwd-qPCR mouse** | CCA AGG ATG GAT ACG TGG AA |
| **Hspb8 rev-qPCR mouse** | TTG GTG AAG TTC TTG GAG ACA AT |
| **Hspb7 fwd-qpCR mouse** | TGC CTA CGA GTT TAC AGT GGA C |
| **Hspb7 rev-qpCR mouse** | TTC ATG ACT GTG CCA TCA GC |

qPCR primers
